# Cryopreservation of Juvenile *Mytilus galloprovincialis*: Safeguarding Mollusk Biodiversity and Aquaculture

**DOI:** 10.1101/2025.01.08.631864

**Authors:** A. Lago, J. Troncoso, E. Paredes

**Affiliations:** Centro de Investigación Mariña (CIM), Departamento de Ecoloxía e Bioloxía Animal, Grupo ECOCOST, Universidade de Vigo

## Abstract

*Mytilus galloprovincialis* Lamarck, 1819, is a globally significant aquaculture species, contributing 25% of fresh seafood landings and leading EU mussel production. In Spain, it accounted for 255,218 tonnes in 2022, around 60% of the global harvest. This study introduces groundbreaking cryopreservation, being the first to successfully cryopreserve the juvenile stage of a marine organism, the largest marine organism ever cryopreserved. By optimizing feeding strategies, equilibration times, and cryoprotectant concentrations, we extended our initial protocol for young larvae to include complex larval stages (72h–26 days post-fertilization) and juveniles (40–45 days, up to 1 mm). These protocols mark a significant step in biodiversity conservation, seedstock preservation, genetic line maintenance, and sustainability of overexploited mollusk populations.

## Main Text

*Mytilus galloprovincialis* Lamarck, 1819, is among the most extensively farmed species globally, accounting for 25% of fresh seafood landings. In recent years, it has become the leading cultivated species in the EU ^1,2^In Spain, specifically, mussel production reached 255,218 tonnes in 2022, representing approximately 60% of the global harvest. Beyond its economic significance, mussel farming has profound sociocultural significance, particularly in regions such as Galicia, where it is integral to culture, the landscape and lifestyle, providing substantial direct and indirect employment. Mussel cultivation typically starts with seed collection sourced from natural beds, which contribute approximately 60–70% of the total, or harvesting from collector ropes suspended from rafts (30–40%). Farmers then attach the seeds to cultivation ropes by hand or with machines secured with rayon or cotton mesh. In NW Spain, mussels generally reach commercial size after 8–13 months. However, this growth rate varies with environmental conditions and location^3^.

In recent years, its cultivation has faced challenges, including declining natural recruitment events and regional restrictions on natural seed collection areas, both of which are critical for traditional raft-based farming. EU mussel production peaked at 600,000 tons in 1990 but declined by 20% to 480,000 tons in 2016 due to factors such as recruitment problems, diseases, algal blooms and other challenges, such as rising sea temperatures, climate change, and/or exposure to pollutants ^4–7^. This study presents flexible cryopreservation protocols for *M. galloprovincialis* larvae and juveniles as a viable tool for aquaculture. This protocol enables the preservation of excess larvae and seed stocks during peak years, maintains valuable genetic lines, and reduces the dependence on seasonal recruitment. Additionally, it would support the restoration of overexploited natural populations ^1,8^.

We developed a successful cryopreservation protocol for mussel larvae at 24–72 hours post-fertilization, facilitating long-term studies on the survival and development of cryopreserved organisms across generations ^9^. On the basis of these findings, we report the effects of feeding in the 48 hours prior to cryopreservation at 72 hours post-fertilization (hpf), modification of equilibration times and adjustment of cryoprotectant concentrations. These results led to the formulation of an advanced protocol for the cryopreservation of significantly more complex larval stages from 72 hours to 26 days post-fertilization (dpf) and even juveniles at 40 and 45 days (Fig. 1) up to 1 mm in size.

**Figure 1.**
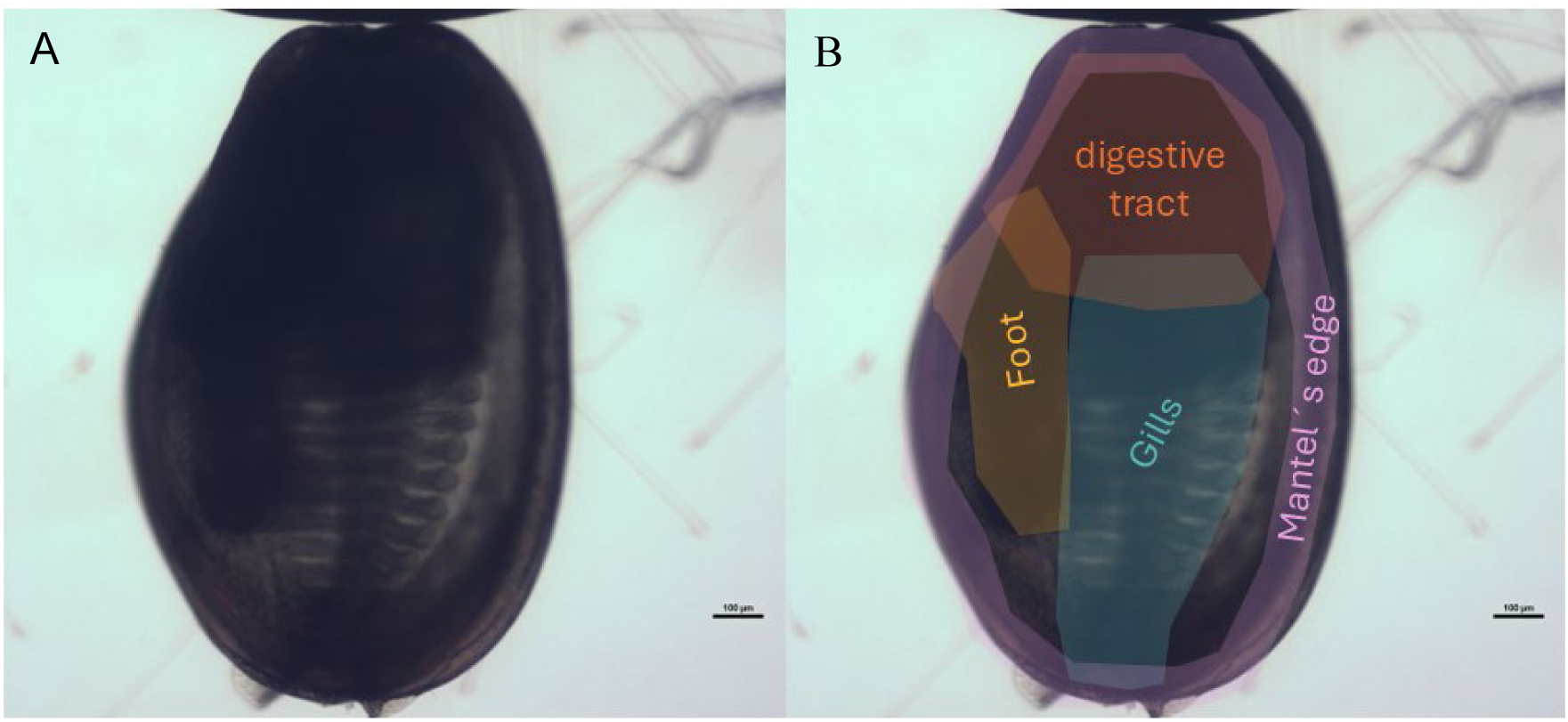
Image showing a 45-day post-fertilization (dpf) juvenile of M*. galloprovincialis* in a general view (A). In (B), the 45-dpf juveniles are depicted with highlighted areas: mantle edge (pink), foot (yellow), digestive tract (orange), and gills (blue).

## Results

### Increasing larval fitness prior to cryopreservation increases the efficiency of the current protocol by 10%

The D-larvae of *M. galloprovincialis* from adult specimens in the Vigo estuary withstand a decrease in salinity of approximately 14.29% and an increase of 21.42% during the first 72 hours of development. Based on the results of size and % normality, we can conclude that the optimum salinity range for larval development is between 30-42.5‰ (Fig. S1A-B). Previously, ^10^ and His et al., 1989^11^ determined that the optimum salinity is 35‰. However, the effects of salinities higher than 35‰ were not analyzed, as presented in this work. The wide range of salinities determined to be suitable for larvae opens the possibility to use beneficial pre-freezing higher salinities^12^, but this approach is not feasible at large aquaculture facilities unless they are situated in regions where the natural salinity is equally high. The cultures at various temperatures (from 12 to 25°C) revealed that the optimal temperature range for the development of D-larvae at 72h of life was between 16-20°C (Fig. S2A-B), with 16°C used as our control and the usual standard procedure. Our temperature range data disagrees with those obtained by Lazo & Pita, 2012^13^, who determined that the optimal temperature range for the growth of *M. galloprovincialis* larvae from the region of Galicia is between 20-24°C. However, our data revealed a large decrease in the percentage of normality at 25°C, which was not as evident in the size of the larvae, although there were significant differences with respect to the control at 16°C. With these results, we decided to increase the incubation temperature standard from 16°C to 20°C for the next experiments to ensure high-quality larvae.

Mussel larvae start feeding at 48 hours when they reach the D-larval stage. In our prior publications, the larvae were cryopreserved at 48 hours and therefore not fed prior^14^. When our data revealed that older larvae were more resistant to cryopreservation, the protocols were modified to use 72h larvae, and we continued without feeding, as a full stomach could be a source of internal ice formation during freezing in such complex larvae. The results of the cryopreservation of 72h-old larvae after feeding (Fig. 2A) show significant differences in larval size between the controls, and these differences can also be observed between the cryopreserved larvae with and without food. The percentage of larval normality significantly differed between fed (84.3±0.9%) and unfed larvae (74.4±5.4%) in both the control and cryopreserved larvae. On the other hand, no significant differences were detected between the fed control larvae and the fed cryopreserved larvae, contrary to what happens if we compare the unfed cryopreserved larvae with their nonfeed control (Fig. 2B).

**Figure 2.**
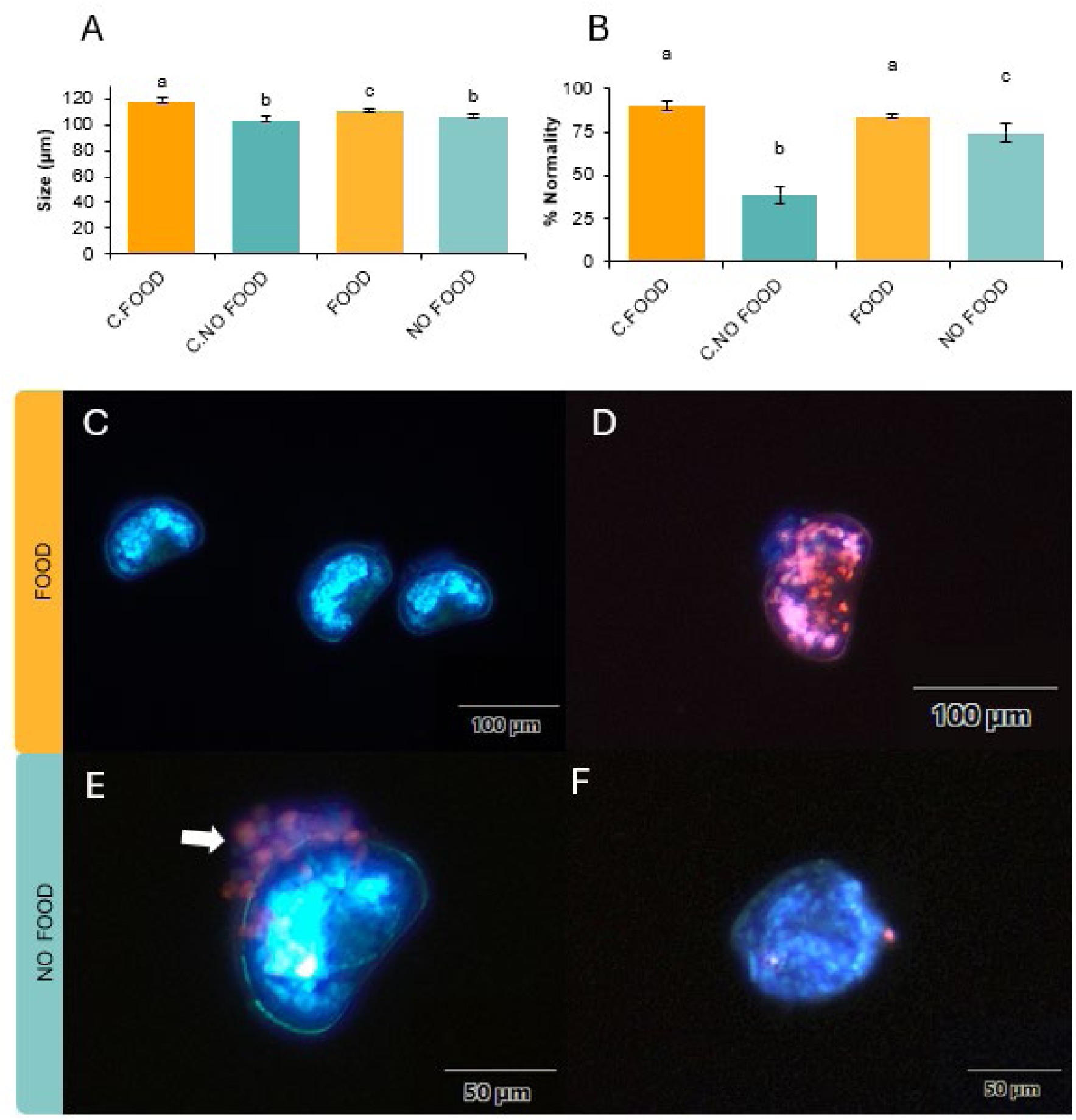
Graph of the mean growth ±SD of cryopreserved larvae at 72 hpf and cryopreserved unfed larvae at 72 hpf at 48 h post-thawing (A). For each parameter, 35 larvae were measured (n=3). Graph of the percentage of larval normality ±SD of cryopreserved feed-fed larvae at 72 hpf and cryopreserved unfed larvae at 72 hpf 48 h post-thawing. (B) For each parameter, 100 larvae were examined (n=3). Asterisks represent significant differences compared with the control without cryoprotectant; * indicates p≤0.05. Images of cryopreserved fed (C and D) and unfed (E and F) larvae of *M. galloprovincilis* at 48 hours post-thaw (hpt), stained with propidium iodide (red) and Hoechst (blue). Larvae cryopreserved after feeding, showing no damage and minimal variation between larval stages (C). Negative control (D). Unfed cryopreserved larvae showing localized damage in the ciliary veil area (thick arrow) (E) and some variation between larval stages (trochophore with slight damage) (F).

We succeeded in cryopreserving, for the first time, D-larvae of *M. galloprovincialis* 72 hpf after feeding. Moreover, all fed larvae, whether cryopreserved or not, were greater in size and percentage of normality than unfed larvae. In the case of the fed cryopreserved larvae, the average size was slightly larger (111±1.64 µm) than that of those cryopreserved without food (107 ±1.61 µm). In the case of % normality, the difference was much greater, ranging from 84.3%±0.9% in fed cryopreserved larvae to 74.4%±5.4% in nonfeed cryopreserved larvae. This implies that feeding the larvae improved their overall fitness, increasing their survival after thawing by approximately 10%.

When the percentage of larval normality of our nonfeed cryopreserved larvae (74.4%) was compared with the current protocol, this percentage was close to that obtained by Heres, 2021^1^ (77%± 4.31%). The survival of cryopreserved larvae after feeding increased by 7.3% (84.3±0.96%).

To obtain the fittest larvae prior to starting the cryopreservation process, we decided to culture them at 35-37%, at 20°C and with a microalgal diet (section 3.6).

### Damage analysis of cryopreserved D-larvae 72 h after feeding reveals decreased damage and improved development synchronicity

The results of the Hoechst and propidium iodide fluorescence dyes revealed small differences in the damage produced between fed and unfed cryopreserved larvae, which were located mainly in the ciliated part of the veil (Fig. 2E). Larvae cryopreserved after feeding do not present this damage (Fig. 2C), which appeared sporadically in unfed larvae. In addition, for larvae that were fed prior to cryopreservation, the divergence between larval stages was minimal, whereas this high variability phenomenon was high in the case of unfed larvae, with trochophore larvae (a prior development stage that should have been overcome before 20 hpf) appearing sporadically (Fig. 2F). To date, no other records of larvae or organisms have been cryopreserved after feeding. Moreover, few previous studies have analyzed the effects of pre-freezing diets after successful cryopreservation. Some works have used different pre-freezing diets on parental lines to modify the lipid composition of sperm, embryos, human T cells, etc. ^15–17^. However, in Guardieiro et al., 2014^16^, the cryotolerance of beef heifer embryos was compromised when pre-freeze diets were used. The same is true for boar sperm fed DHA-enriched diets^17^. In contrast, in the case of Baynes & Scott, 1987^15^, the use of pre-freeze diets increased viability and the post-thaw proliferation rate in human T cells. If we analyze our results together with those of the few previous studies, we can assume that it is not so much the use of lipid-enriched diets that is important but rather the diet itself.

The lower damage in fed larvae disproves the assumption that the full stomach could be a source of ice formation but also shows that larvae cryopreserved after feeding are greater in size and have higher survival rates. In contrast, the commonly reported developmental delays and lesions associated with cryopreservation^9,18^ are reduced. Our results suggest that feeding larvae before freezing significantly enhances both their survival and size, providing additional protection against cryopreservation, an important factor to consider when optimizing existing protocols.

### Cryopreservation of larvae throughout their larval rearing cycle as a tool for aquaculture management and biodiversity conservation

By feeding larvae prior to cryopreservation, it was possible to study the success of cryopreserved larvae as their development progressed all the way to their settlement and transformation into juveniles. We successfully cryopreserved larvae at 24 hpf, 48 hpf, 72 hpf, 144 hpf, 10 dpf (10d), 15 dpf (15d) and 20 dpf (20d), as well as juveniles at 40 dpf (40d*) (Figure 3). These achievements marked a significant milestone in cryopreservation research and paved the way for more intricate studies into long-term viability, growth patterns, and potential applications in aquaculture and conservation biology.

**Figure 3.**
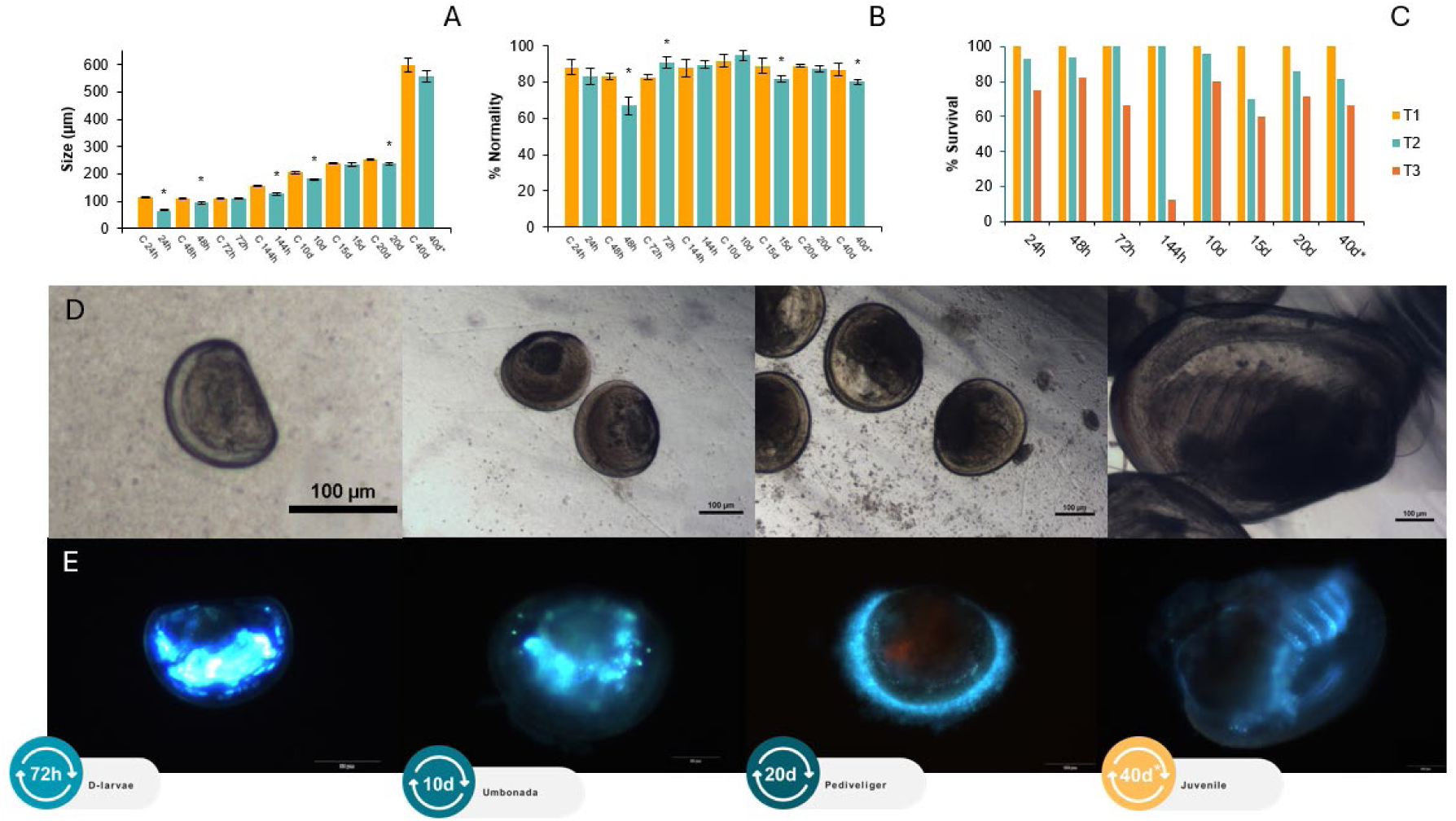
Graph of the mean growth ± SD of cryopreserved larvae and control larvae throughout larval development from 24 hpf until 40 dpf * (40d * last modification protocol), 48 h post-thawing (A). For each parameter, 35 larvae were measured (n=3). Graph of the percentage of larval normality ± SD of cryopreserved larvae and their controls throughout larval development from 24 hpf until 40 dpf * (40d* last modification protocol), 48 h post-thawing (B). For each parameter, 100 larvae were examined (n=3). Asterisks represent significant differences compared with the control without cryoprotectant; * indicates p≤0.05. Images of control larvae obtained via light microscopy (D) and images of cryopreserved larvae after 72 hours stained with propidium iodide and Hoechst, obtained via fluorescence microscopy, along different stages of larval development of *M. galloprovincialis*.

The size (Fig. 3A) and percentage of normality (Fig. 3B) of cryopreserved larvae throughout all developmental stages was smaller than that of the controls, similar to the work of Rusk et al., 2020^18^ (Fig. 3A). However, these differences are small and seem to disappear as larval development progresses. It is possible that at times of intense development, changes in the cryopreservation process further accentuate the metabolic expenditure necessary for normal development. Moreover, cooling and thawing cryoinjuries could be more critical in each specific development stage. This results in slight delays in larval growth, but as indicated in Heres et al., 2022^9^ and Paredes *et al*, 2015^19^, this delay is minimized with the passage of days post-freezing.

### Damage analysis of serial cryopreservation of larvae

Immediately after thawing (T1), larvae from all the studied ages were alive, with no dead or damaged larvae observed in the samples. However, the survival analysis indicated a decline in survival percentage over time. The percentage of larvae surviving 48 h post-thaw (T2) exceeded 60% in all cases, except for the 144hpf larvae. The low survival rates in these developmental stages (approximately 13%) may be due to different causes (Fig. 3C). The reduced survival of 144 hpf larvae could be attributed to the larvae being in a sensitive developmental phase during cryopreservation, such as during the formation of critical tissues or organs or the transition to a new larval stage. In our case, when the larvae were incubated at 20°C, at 144 h, our larvae had already left the D-larvae stage and started to transform into umbonate at this time, and the gill septa and the rudiment of the foot began to form^20^. Therefore, damage to these structures or metabolic energy expenditure due to cryopreservation may be behind this decrease in the post-thaw survival rate.

However, it is important to note that the analysis was conducted without replicates, focusing solely on the total number of larvae present in 1 ml of each sample.

Using fluorescence staining (Fig. 3D), we identified common lesions of larvae and juveniles, which were predominantly localized in the digestive region. Although propidium iodide staining usually marks this region in general, even in controls, we occasionally observe small vesicles, some of which appeared to exhibit movement in the region; therefore, we suspect that these vesicles may have been remaining or living algae into which the dye may have penetrated when the cell membranes were compromised during the digestive process, i.e., half-digested microalgae. In any case, we cannot rule out the possibility that lethal or sublethal cryoinjury may occur in this region of the body, so in the future, it would be advisable to analyze this region in depth in a more specific way via histological, immunohistochemical or scanning microscopy techniques.

On the other hand, all the data presented from the size, survival and post-thaw damage analyses of 40d* juveniles are the result of the latest specialized modifications of the cryopreservation protocol. Details of the experiments performed to develop these modifications are provided in the following sections.

### How to increase survival in cryopreserved juveniles

Initial attempts to cryopreserve 40-day-old juveniles (40 dpf) via the protocol developed by Heres (2021)^21^ were successful. However, analysis of survival rates over 48 hours post-thaw revealed a significant decline, with survival decreasing to approximately 13% at 48 hours (*T3*) (Fig. 4B). These juveniles, which are substantially larger (approximately 500 µm) than 72-hour D-larvae (approximately 100 µm) and possess fully developed organs and tissues such as adults, may require adjustments to the cryoprotectant concentrations. Specifically, the concentration used in the current protocol used for the larval stages (10% ethylene glycol [EG] + 0.4 M trehalose [TRE]) may be insufficient to provide adequate protection for larger, more complex individuals.

**Figure 4.**
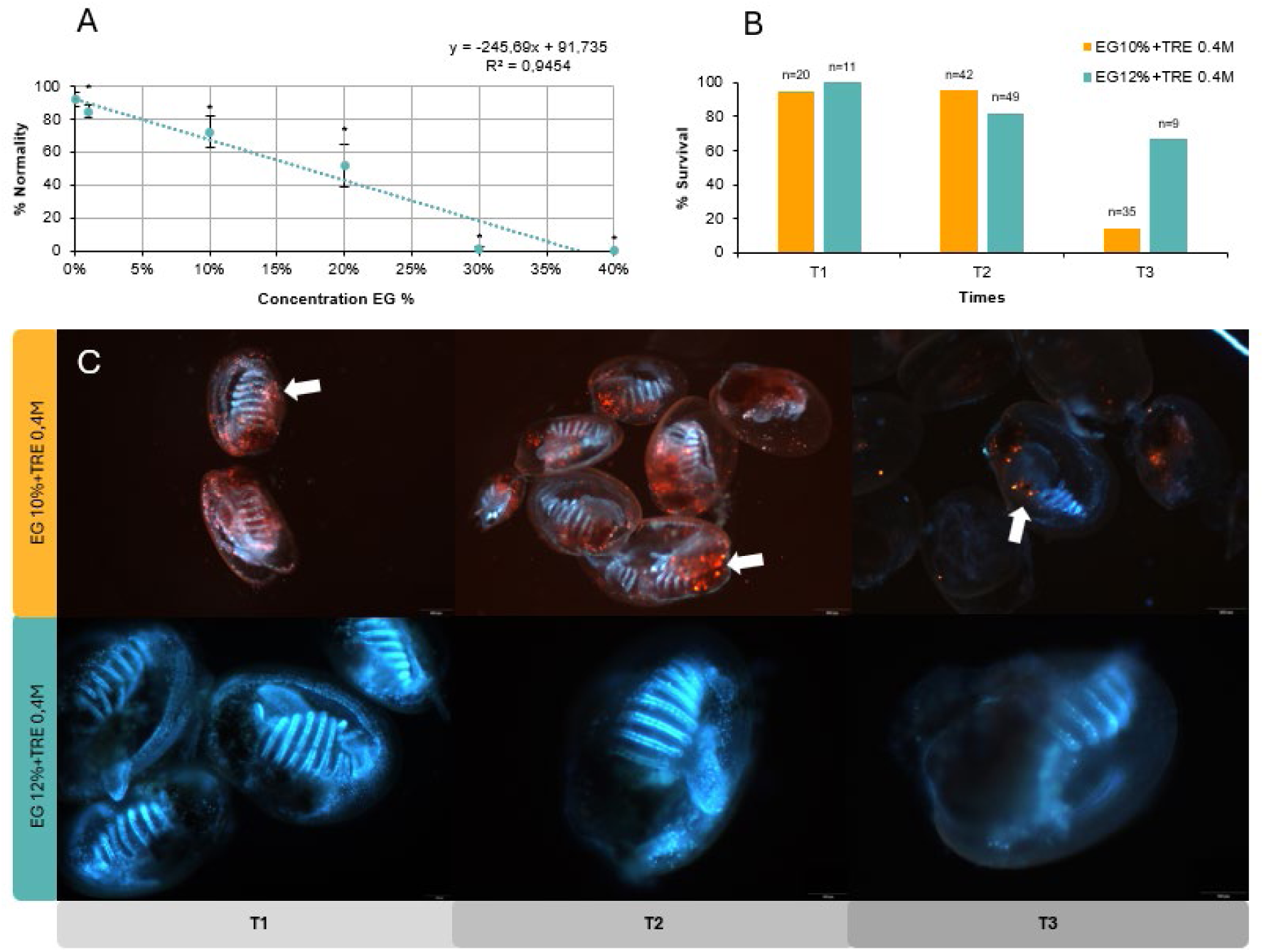
Dose‒response curve of 40-day-old *M. galloprovincialis* juveniles exposed to different concentrations of EG. For each parameter, 100 larvae were measured (n=4) (A). Graph of % survival of cryopreserved juveniles at 40 dpf at three time points, T1 (0 h after thawing), T2 (24 h after thawing) and T3 (48 h after thawing), analyzed via the fluorescent live-dead dye for two concentrations of cryoprotectant, EG10% or 12% with a TRE of 0.4 M, n= number of juveniles analyzed per treatment as present in 1 mL of sample (B). The thick arrow shows particulate material expelled by the juvenile after thawing. Images of cryopreserved juveniles fed with EG10% TRE 0.4M and of cryopreserved juveniles fed EG10% TRE 0.4M at three time points, T1 (0 hpt), T2 (24 hpt) and T3 (48 hpt), with propidium iodide and Hoechst dye (C).

The toxicity of ethylene glycol was studied through a dose‒response curve (Fig. 4A). The results revealed significant differences from the control at all EG concentrations. As the EG concentration increased, the survival rate decreased, reaching zero at the highest EG concentration (40% or 7.17 M EG). From the equation of this curve, we calculated the IC50 of EG specifically for 40 dpf juveniles, which corresponds to 16.8% EG (3.03 M). Since our goal was for juvenile survival to be greater than 50%, we decided to increase the EG concentration from 10% (1.79 M) to 12% (2.15 M) for juveniles. With this adjustment, we aim to increase the level of cryoprotection of our juveniles without significantly increasing toxicity, which could compromise their survival.

### The cryopreservation of 40 dpf juveniles requires a relatively high cryoprotectant concentration

The survival percentage results are shown on Fig. 4B. An increase of approximately 52% at time T3 (48 h post-thawing) with respect to the juveniles cryopreserved with EG 10%+TRE 0.4M TRE at this time. With this new cocktail, we increased the survival rate from approx. 13% to approx. 66% at 48 h post-thawing. It is evident that at 24 h post-thawing (T2), the % survival was greater with EG10%+TRE 0.4M than with EG12%+TRE 0.4M, however, at T3, the decrease in % survival was very evident with EG10%+TRE 0.4M (Fig. 4). In 40 dpf juveniles (EG10%+TRE 0.04M), a large amount of particulate matter was concentrated in the digestive tract region (thick arrow). These particles are expelled from the inside of the animal to the outside. Furthermore, these particles are stained with propidium iodide, which indicates that they are dead cells. We do not know the specific origin of these particles, but we suspect that they may be food remains (microalgae) or remains of some region of the digestive tract that is slowly disintegrating. However, once the 40 dpf* (EG10%+TRE 0.04M) juveniles were cryopreserved with the new cocktail, the presence of this particulate material was practically nil (Fig. 4C). This allowed us to successfully cryopreserve 40 dpf juveniles (40d*), the % normality, length and survival data of which are discussed above (Fig. 3). On the other hand, no major differences were observed in % survival between treatments except for T3 (48 h post-thaw), when the increase in cryoprotection seemed to yield an increase in survival, as there were no replicates in this case, and no statistical analysis was performed at this specific point (due to the nature of the sample, as explained in section 3.5).

### Cryopreservation of 45 dpf juveniles with the improved protocol results in 90% survival

Ultimately, we cryopreserved 45-day-old juveniles with EG12%+TRE 0.4M, and this time, we introduced a brief fasting period (96 h). We expected the introduction of fasting to help obtain a better survival rate. The size and percentage of normality data are very favorable at 48 h post-thawing (Fig. 5A, 5B) and was not significantly different from that of the control. In this way, we managed to cryopreserve 45-day-old juveniles with a size of more than 1 mm and maintained a normality percentage of more than 90%.

**Figure 5.**
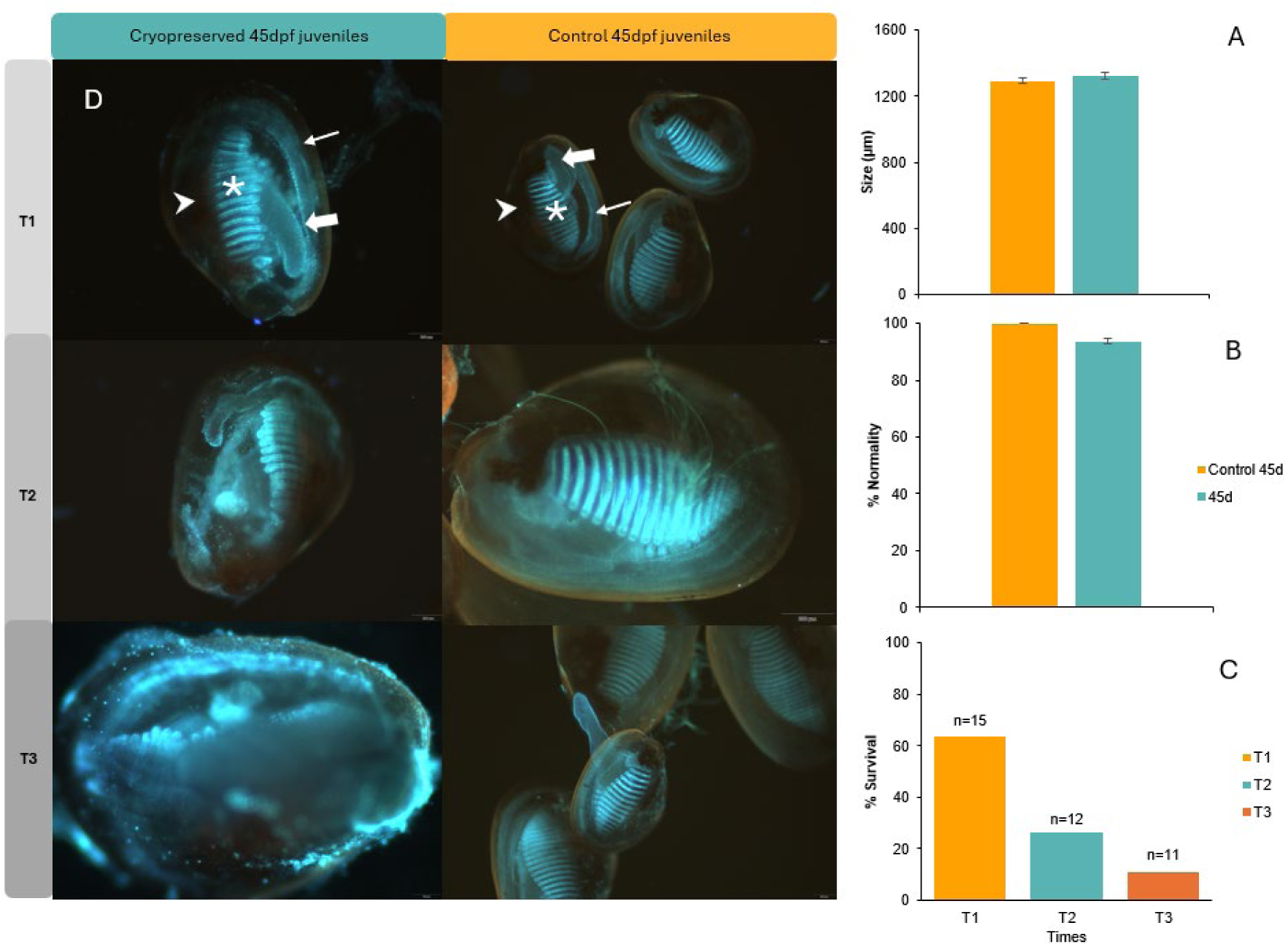
Graph of the mean growth ±SD of cryopreserved feed 45 hpf juvenile and control 45hpf juvenile 48h post-thawing (A). For each parameter, 35 larvae were measured (n=3). Graph of the percentage of larval normality ±SD of cryopreserved feed 45hpf juvenile and control 45hpf juvenile 48h post-thawing (B) For each parameter, 100 larvae were examined (n=3). There were no significant differences (p≤0.05) compared with the control without the cryoprotectant. Graph of % survival of cryopreserved juveniles at 45 dpf cryopreserved with EG 12%+TRE 0.4 M at three time points, T1 (24 h after thawing), T2 (48 h after thawing) and T3 (72 h after thawing), analyzed via the fluorescent live-dead dye EG 12%+TRE 0.4 M, n= number of juveniles analyzed per treatment as present in 1 mL of sample (C). Images of cryopreserved juveniles fed EG12% +TRE 0.4M and of cryopreserved juveniles fed EG12%+TRE 0.4M at three time points, T1 (24 hpt), T2 (48 hpt) and T3 (72 hpt), with propidium iodide (red, dead cells) and Hoechst (blue, active DNA) dye (D). The thin arrow points to the edge of the mantle, the thick arrow to the foot, the caret to the digestive tract and the asterisk to the gills.

### Damage analysis of serial cryopreservation after feeding, new cocktail addition and fasting

The survival rate of 45-day post-fertilization (dpf) juveniles decreases over time following cryopreservation. After the first 24 h post-thawing (T1), the survival rate was approximately 63%, which decreased to approximately 11% by 72 h post-thawing (T3) (Fig. 5C). Although this 72-hour survival rate may seem low, it is important to note that this is the first successful cryopreservation of juvenile mussels, which are 10 times larger than the size for which the current protocol was designed. Moreover, we are dealing with a fully developed organism that is indistinguishable from an adult, except for its size, and presents significant metabolic and systemic complexity. Thus, this represents the cryopreservation of an entire organism. In the images of Fig. 5C. no differences at the structural level were evident between the 45 dpf juveniles subjected to cryopreservation and the control juveniles. The main anatomical structures, such as the gills, foot, mantle edge and digestive region, appeared to remain intact in both treatments. In addition, no areas marked with propidium iodide were detected, suggesting the absence of cell damage in juveniles that survived the cryopreservation process during the first 72 hpt.

After thawing (24 h), a high filtration rate can be observed in the gills, as well as some movement in the foot and mantle (Movies S1, S2). The byssus allows slight fixation, but this can be easily detached. At 48 h post-thawing, filtration and shrinkage of the mantle and foot were less evident than they were initially post-thawing. However, byssus formation is much clearer, and its binding power is more pronounced. We suspect that first, the filtration rate increases because juveniles try to eliminate the remains of the CPAs from their organisms, also with the help of slight contractions of the mantle. Moreover, during the cryopreservation process, partial or total denaturation of the byssus proteins may have occurred because of the cold and exposure to the CPAs. On the other hand, the weakening of the byssus as an attachment structure to the substrate and to the rest of its congeners causes them to use the foot as a way of detecting the proximity of other juveniles in search of protection. Finally, the formation of new byssus strands and the increase in attachment strength indicate that the byssus gland is not damaged during cryopreservation process. In the future, it would be desirable to try to obtain hard evidence to validate this hypothesis quantitatively.

Owing to the technical complexity of juvenile analysis, it was not possible to perform replicates for a more detailed assessment of survival rates via this fluorescence technique. The fact that these organisms have byssus implies an extra difficulty. Naturally, the individuals of this species use the byssus structure to adhere to each other and to surfaces. This means that they appear in small aggregates of several individuals, making it very difficult to separate and work with a specific number of individuals in each straw. Despite this limitation, the preliminary data obtained were consistent across trials and provided valuable insights into the timing and location of lethal and sublethal cryoinjuries in larvae and juveniles. While the lack of replicates limits precise quantification, the consistency observed suggests that the results are representative of the effects of cryopreservation under the current experimental conditions. Moreover, this approach has proven useful as an exploratory tool to detect general patterns of cryodamage, offering critical insights into the processes that influence post-cryopreservation survival. For future studies, the optimization of this methodology is anticipated, along with the potential for additional replicates or alternative approaches to allow for more accurate survival quantification of organisms with a byssus that aggregates and attaches together.

To date, successful cryopreservation of *Mytilus galloprovincialis* larvae beyond 72 hours post-fertilization (hpf) has not been achieved. However, we have not only surpassed this developmental stage but also successfully cryopreserved, for the first time, all larval stages of this species, including juvenile individuals. This makes us the first to cryopreserve all larval stages of any species globally and among the few studies to have successfully cryopreserved juveniles. These juveniles, both metabolically and in terms of tissue and organ complexity, are indistinguishable from adult individuals, except for their size.

Organismal cryopreservation studies are very novel, with few published examples, such as *Caenorhabditis elegans*^22^, which was cryopreserved via isochoric cryopreservation, and survival was tested for up to 24 hours. Coral fragments from the species *Porites compressa*^23^ were successfully cryopreserved to a size of 1 cm. Other complex multicellular systems, such as embryos and larvae, can be equally important and comparable in size. Notable examples include the cryopreservation of *Drosophila melanogaster* embryos^24^, zebrafish *Danio rerio* embryos^25^, and shrimp larvae^26^, all of which are comparable in size to our 40 dpf larvae (more than 1 mm). These studies have successfully addressed the challenge posed by the low surface-to-volume ratio, which reduces the efflux of water and cryoprotectants (CPAs). In the case of mussels, the animal body inside the shell has an elongated and hollow shape, making the penetration of cryoprotectants easier, as the surface-to-volume ratio is larger than the size suggests; on the other hand, their shells are impermeable to water and cryoprotectants, and mussels can open/close at will react to the stimuli of the environment (like when detecting a chemical in the water such as a cryoprotectant) ^27^; therefore, this poses a completely unique set of challenges and advantages that have allowed us to cryopreserve mussel juveniles via slow cooling instead of vitrification.

Working with gregarious organisms poses the challenge of handling their naturally formed aggregates, which they use for protection. These juveniles attach to each other and to surfaces through the byssus, a protein-based structure critical for their survival. During the experiments, it was necessary to carefully detach the settled juveniles from the tanks, create aggregates of consistent numbers and sizes, and work quickly to prevent them from reaching the experimental surfaces. Although cryopreservation may cause byssus denaturation immediately after thawing, the integrity of the byssal gland allows juveniles to regenerate new byssus and restore its normal function.

This research lays an essential foundation for improving cryopreservation techniques and deepening our understanding of biological responses at both the cellular and systemic levels in whole organisms.

## Acknowledgements

This work is part of the Galicia Marine Science program in Complementary Science Plans for Marine Science of Ministerio de Ciencia, Innovación y Universidades included in the Recovery, Transformation and Resilience Plan (PRTR-C17.I1).

## Author contributions

Conceptualization: AL, EP, JT

Methodology: AL

Investigation: AL

Visualization: AL, EP

Funding acquisition: EP

Project administration: EP

Supervision: EP

Writing – original draft: AL, EP

Writing – review & editing: AL, EP, JT

## Data and materials availability

The authors are submitting the raw data to (Integrated Marine Information System) IMIS, and we will include the DOI of the data in this manuscript before publication.

## Supplementary Materials

### Materials and Methods

#### 1.1. Larvae collection

Mussels (*M. galloprovincialis*) were provided by the biological resource service at Estación de Ciencias Mariñas de Toralla (ECIMAT) of the Centro de Investigación Mariña (CIM)-Universidade de Vigo. Gametes were obtained via thermal cycling. In the case of the feeding cryopreservation experiment, a single pair of males and females was used to minimize the noise that genetic variation can add to the assay. Finally, in the rest of the experiments, owing to the high requirement of genetic material, a pool of three males and three females was used. The quality of both spermatozoa and oocytes (sperm motility, oocyte sphericity and color homogeneity) was evaluated. After a contact period of 15 min, the percentage of fertilization was assessed via visual verification of the formation of the polar corpuscle under a light microscope (Nikon Eclipse TE2000). The fertilization rate in all the experiments was greater than 80%.

#### 1.2. Cryopreservation methods

Our protocol builds on that developed by Heres, 2021^21^, in our laboratory; we used a Freeze Control System Cryologic Pty Ltd., Mt Waverley, Australia). The protocol consisted of cooling from 4°C to -35°C at a rate of 1°C/min, with a 2 min pause at -12°C, at which time seeding was checked and induced, if necessary, in the straws. Cryopreservation experiments were performed in 0.25 ml straws, except for the cryopreservation of 40 dpf and 45 dpf juveniles, where 0.5 ml straws were used. The punctual use of the 0.5 ml straws was due to the size of the juveniles, whose diameter was similar and even sometimes larger than that of the 0.25 ml straws. Thawing of the straws was done by immersion in a water bath at 35°C for 6 sec, 8 sec in the case of the larger straws. Finally, dilution and addition of the CPAs occurred in a single step (1:1) in FSW (1 µM +UVA) at room temperature (19±1°C).

The equilibration time used was 15 min for the cryopreservation of larvae up to 48 hpf, which was subsequently increased to 1 h according to Heres *et al*. (2021)^21^, allowing for the equilibration of larger larvae with a shell. The concentration of CPAs was modified as explained in section 3.6.

#### 1.3. Incubation of 72-h-old D-larvae for cryopreservation after feeding

After fertilization, the larvae were incubated at 20°C in an incubator in a 2 L glass beaker with 300 larvae/ml UV-FSW (1 µm) with light aeration until they reached 72 h of life. Larvae were fed immediately after reaching the D-larvae stage at 48 h with a bialgal diet consisting of 40 equivalents of *Chaetoceros neogracilis* VanLand, 1968, and *Tisochrysis lutea* El M. Bendif & I. Probert, 2013. After 72 h, the larvae were cryopreserved following the current protocol for mussel cryopreservation (Heres, 2021). The control was incubated and handled under the same conditions as the other larvae and was not fed for the first 72 h of life. Afterward, the control was fed the same diet as described above alongside the thawed larvae.

#### 1.4. Incubation of larvae and juveniles for serial cryopreservation with feeding

Five million embryos were incubated at 20°C in an isothermal room in 100 L tanks with UV-FSW (1 µm) and light aeration. Larvae were fed after reaching the D-larvae stage at 48 h of life *ad libitum* with five algae diets, *C. neogracilis*, *T. lutea, Rhodomonas lens* Pascher & Rutter, 1913*, Phaeodactilum tricornutum* Bohlin, 1897 *and Tetraselmis suecica* Butcher, 1959 (food amounts increased according to larval stage requirements). A subsample of larvae was then filtered, concentrated and cryopreserved at different developmental stages during the 20 days of larval rearing.

#### 1.5. Salinity and temperature bioassays

The effects of exposure to 9 different salinities (20‰, 22.5‰, 25‰, 30‰, 35‰, 37.5‰, 40‰, 42.5‰ and 45‰) were evaluated, with 16°C and 35% salinity used as controls (n=4). The effects of temperature exposure were evaluated at 6 different temperatures (4°C, 10°C, 16°C, 20°C, 25°C and 30°C), with 16°C and 35% salinity used as controls (n=4). In both cases, after fertilization, 300 larvae were placed in closed 20 mL vials with ASW (artificial seawater, following the recipe from Lorenzo et al., 2002^28^, which is based on a formulation modified from Zaroogian, 1969^29^) at a controlled temperature and incubated for growth for 48 h under the different conditions described above. Finally, the effects of both parameters on the larvae were evaluated on the basis of the percentage of normality and larval size after 48 h^30^.

#### 1.6. Damage analysis in cryopreserved larvae after feeding

Live-dead fluorescence dye consisting of Propidium iodide (1 mg/ml; staining shows dead cells in red), Hoechst 1 M (live cells with active DNA are shown in blue), and an incubation time of 2 min (adapted from Lema *et al*., 2011^31^) were used. Larval damage was analyzed via Olympus CX43 fluorescence microscopy (CL-CX43-RFA-T). In total, 1 ml of larvae per sample was analyzed.

A negative control of the D-larvae was performed to ensure that the propidium iodide worked perfectly and stained those areas where the cells were dead. For this purpose, several subjects were sacrificed, and the efficacy of the dye was tested (Fig. 2D).

#### 1.7. Toxicity test of the CPAs

A dose‒response toxicity assay for ethylene glycol (EG) was conducted to determine its IC50 value and optimize its concentration as a cryoprotectant. The inhibitory concentration 50 (IC₅₀) is a widely used parameter in toxicology to measure the effectiveness of a substance in inhibiting a biological process. In our case, we used the IC₅₀ in relation to the percentage of normality, defining it as the concentration of a substance required to reduce normality by 50%. The concentration corresponding to the IC50 was 16.9%. Since we wanted to increase the concentration of EG as much as possible and therefore protect the CPA without compromising the % normality, we decided to increase the percentage of EG by only 2% with respect to the original protocol, ranging from 10% EG to 12%. In this way, we ensured normality higher than 50% in our experiments. The IC₅₀ of EG for 40 dpf juveniles was calculated from the equation of the curve resulting from the dose‒response assay of the 5 different cocktails. We prepared five CPAs cocktail solutions (EG 1% + TRE 0.4 M, EG 10% + TRE 0.4 M, EG 20% + TRE 0.4 M, EG 30% + TRE 0.4 M, and EG 40% + TRE 0.4 M). The toxicity assays were carried out in 2 ml cryovials, each containing 40 dpf juveniles of *M. galloprovincialis*. Because juvenile mussels naturally form aggregates by joining at the byssus, an approximate number of juveniles are added in the form of small aggregates, ensuring that all replicates have the same number of individuals. The dilution and addition of the CPAs occurred in a single step (1:1) at room temperature (19±1°C), and the juveniles were exposed for 1 hour. After the exposure period, the juveniles were filtered through 40 µm mesh, and the cryoprotectant residues were removed by rinsing with UV-FSW (1 µm). Finally, the juveniles were transferred to 20 ml bioassay vials containing FSW and incubated for 48 hours in an incubator at 20°C before being fixed with 37% formaldehyde. From the equation of this curve, we calculated the IC50 of EG for 40 dpf juveniles.

#### 1.8. Biometric analysis

Larval and juvenile fitness was determined as follows: growth capacity (n=35) and normal development (n=100) following the criteria developed by Heres, 2021^21^, His et al., 1997^11^, Paredes et al., 2013^32^, Rusk, 2012^33^ and Ventura et al., 2016^34^. These parameters were assessed by calculating the percentage of normality and length (µm) by measuring the maximum length along the anterior/posterior axis parallel to the hinge. Larvae and juveniles with malformations, severely undeveloped margins or hinge deformations (convex or concave hinge), fertilized eggs or protruding mantle were considered nonnormal larvae. In addition, photographs and videos were acquired via a Nikon TE2000S inverted microscope with image analysis (NIS type).

#### 1.9. Data processing and statistical analysis

Statistical analysis of the data for both percentage normality and larval and juvenile size for all bioassays and experiments was performed via SPSS® version 15.0 software. The Shapiro‒ Wilk test or the Kolmogorov‒Smirnov test (depending on the requirement of the data) was used to analyze the normal distribution of the data. For those presenting a normal distribution, Levene’s test for homogeneity was used, and then one-way analysis of variance (ANOVA) was performed, followed by Bonferrony or Tahame, depending on whether the variances were assumed equal or not, respectively. In cases where the data did not fit a normal distribution, a Kruskal‒Wallis test was performed^35,36^.

---

**Fig. S1.**
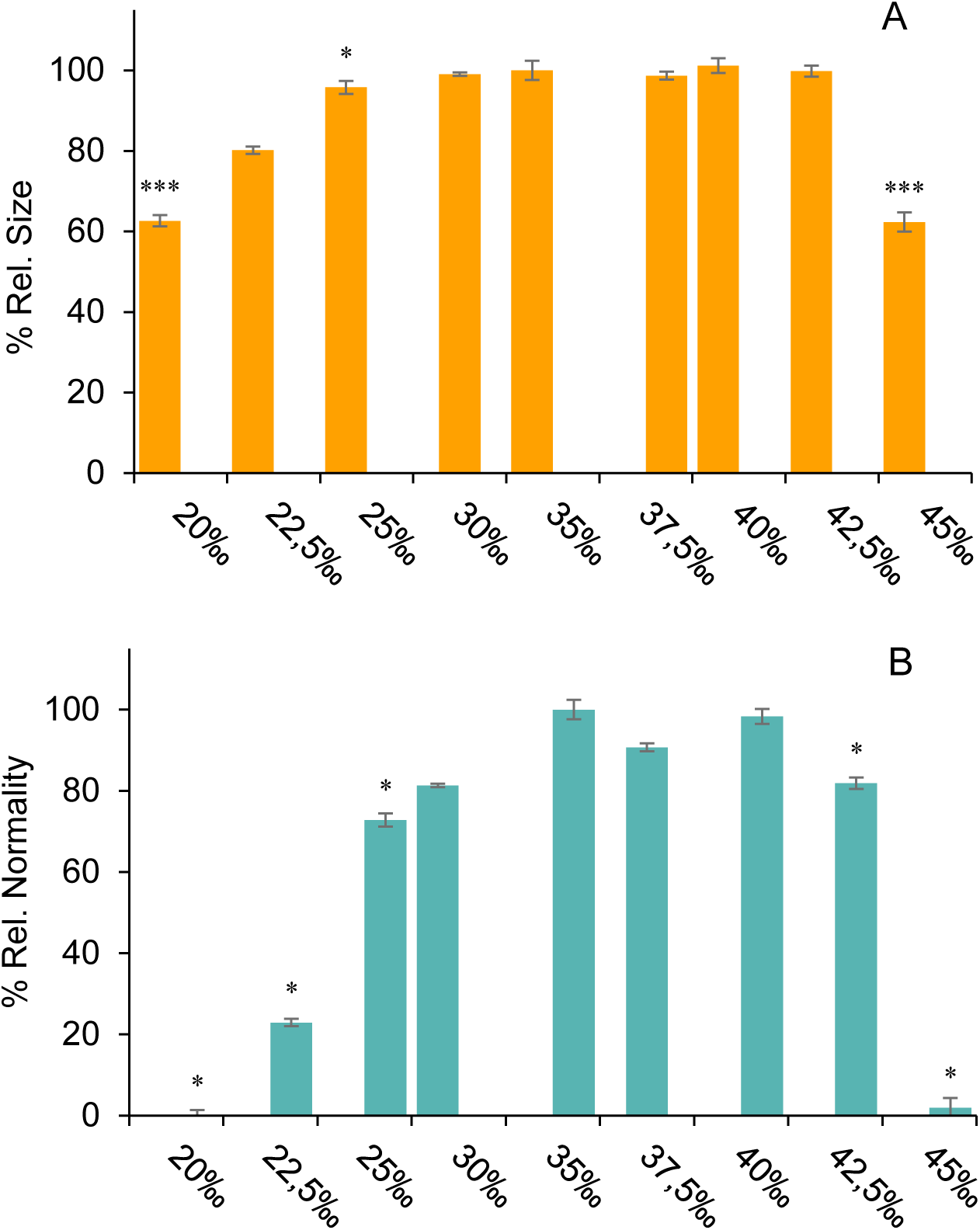
Graph means larvae growth ± SD, versus salinity (A). For each treatment, n=35 larvae were measured, and the experiment was conducted in triplicate. Graph mean percentage of larval normality ± SD, versus salinity (B). For each treatment, n=100 larvae were examined, and the experiment was conducted in triplicate. Asterisks represent significant differences with the control without cryoprotectant ((***) is p≤0.001), (**) p≤0.01) and (*) p≤0.05).

**Fig. S2.**
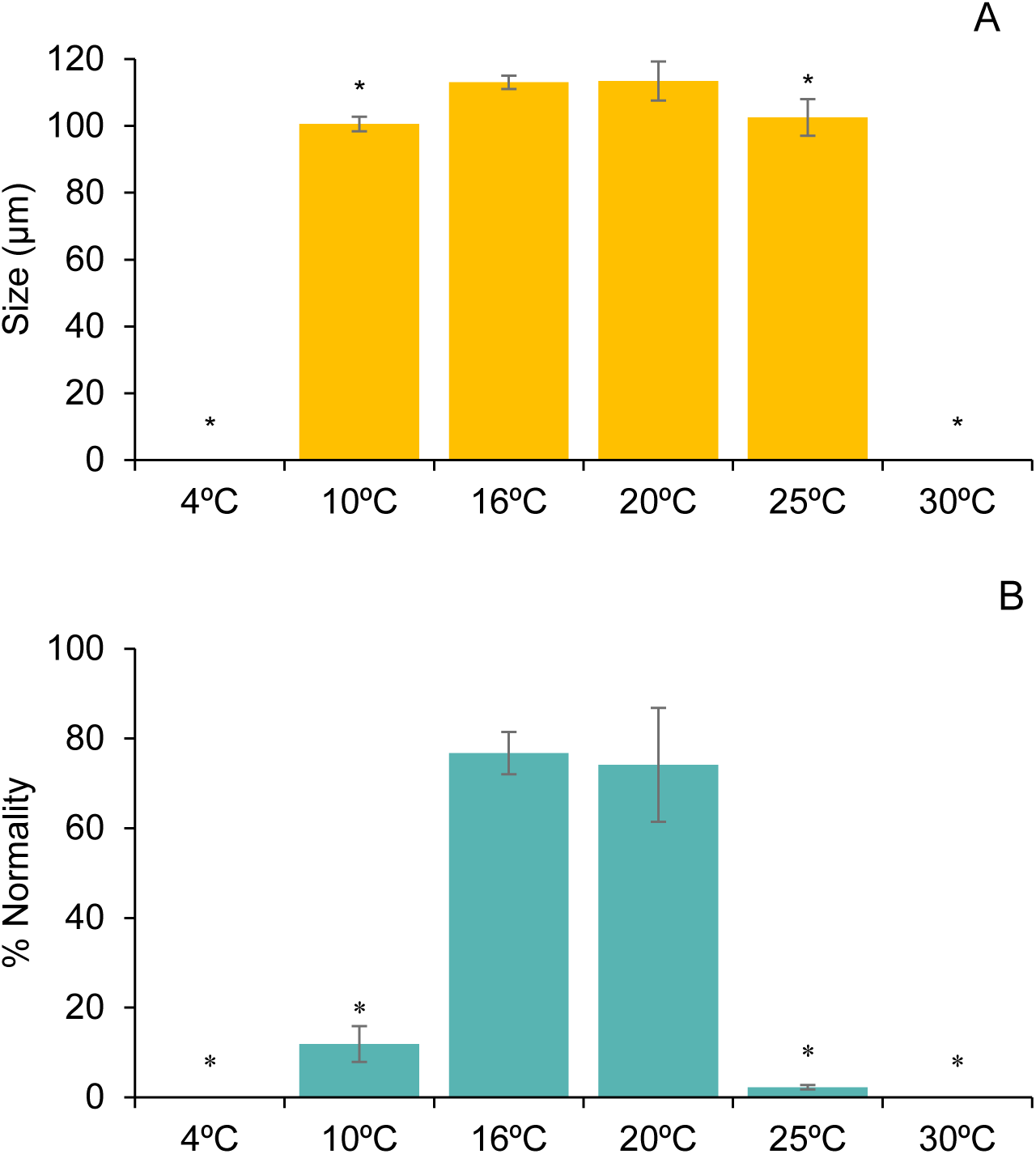
Graph the mean larval growth ± SD, versus temperature (A). For each treatment, n=35 larvae were measured, and the experiment was performed in triplicate. Graph mean percentage of larval normality ± SD, versus salinity (B). For each treatment, n=100 larvae were examined, and the experiment was conducted in triplicate. Asterisks represent significant differences with the control without cryoprotectant ((***) is p≤0.001), (**) p≤0.01) and (*) p≤0.05).

**Movie S1. [click here](https://youtu.be/OPKW6gCDrRs)**

Juvenile 40dpf, 24hpt cryopreserved with EG12%+TRE0.4M.

**Movie S2. [click here](https://youtu.be/ZsWjkhnsN78)**

Juvenile 45dpf, 24hpt cryopreserved with EG12%+TRE0.4M.

